# An inducible ESCRT-III inhibition tool to control HIV-1 budding

**DOI:** 10.1101/2023.10.16.562494

**Authors:** Haiyan Wang, Benoit Gallet, Christine Moriscot, Mylène Pezet, Christine Chatellard, Jean-Philippe Kleman, Heinrich Göttlinger, Winfried Weissenhorn, Cécile Boscheron

## Abstract

HIV-1 budding as well as many other cellular processes require the Endosomal Sorting Complex Required for Transport (ESCRT) machinery. Understanding the architecture of the native ESCRT-III complex at HIV-1 budding sites is limited due to spatial resolution and transient ESCRT-III recruitment. Here, we developed a drug-inducible transient HIV-1 budding inhibitory tool to enhance the ESCRT-III lifetime at budding sites. We generated auto-cleavable CHMP2A, CHMP3, and CHMP4B fusion proteins with the hepatitis C virus NS3 protease. We characterized the CHMP-NS3 fusion proteins in the absence and presence of protease inhibitor Glecaprevir with regard to expression, stability, localization and HIV-1 Gag VLP budding. Immunoblotting experiments revealed rapid and stable accumulation of CHMP-NS3 fusion proteins with variable modification of Gag VLP budding upon drug administration. Notably, CHMP2A-NS3 and CHMP4B-NS3 fusion proteins substantially decrease VLP release while CHMP3-NS3 exerted a minor effect and synergized with CHMP2A-NS3. Localization studies demonstrated the re-localization of CHMP-NS3 fusion proteins to the plasma membrane, endosomes, and Gag VLP budding sites. Through the combined use of transmission electron microscopy and video-microscopy, we unveiled drug-dependent accumulation of CHMP2A-NS3 and CHMP4B-NS3, causing a delay in HIV-1 Gag-VLP release. Our findings provide novel insight into the functional consequences of inhibiting ESCRT-III during HIV-1 budding and establish new tools to decipher the role of ESCRT-III at HIV-1 budding sites and other ESCRT-catalyzed cellular processes.

## 1 Introduction

HIV-1 assembly and budding take place at the plasma membrane [1, 2] and require the interplay of viral structural proteins [3] and cellular factors to release new infectious virions [4-8]. Notably, budding is mostly driven by the polyprotein Gag and its expression suffices to produce virus-like particles (VLPs)[9]. The p6 domain of Gag contains essential late domains [10-13] that have been shown to recruit the cellular endosomal complex required for transport (ESCRT) machinery via Tsg101, a subunit of ESCRT-I [14-16] and/or via the ESCRT-associated factor Alix [17-20].

The ESCRT machinery is composed of five complexes, ESCRT-0, -I, -II, -III and VPS4 [21], which catalyze numerous membrane remodeling processes including topologically similar in-side out budding implicating ESCRT-III and VPS4 in the final cut via membrane fission [22-25]. Mammalian cells express eight different ESCRT-III proteins, CHMP1 to 8, with CHMP1, CHMP2 and CHMP4 present in two and three isoforms, respectively [23]. ESCRT-III proteins shuttle between an inactive autoinhibited closed conformation [26, 27] and an activated state [28-30] that polymerizes in the open ESCRT-III conformation [31-34] assembling into loose filaments or tight helical structures *in vitro* [27, 35-40] and *in vivo* [41-45]. Most ESCRT-III proteins contain MIT-domain interacting motifs (MIMs) present within their C-termini recruiting VPS4 [46]. VPS4 was suggested to constantly remodel ESCRT-III *in vivo* [47] consistent with VPS4-catalyzed ESCRT-III filament remodeling *in vitro* [48, 49]. Notably CHMP2A-CHMP3 filament remodeling into dome-like end-caps was suggested to constrict membrane necks from fixed diameters of 45 to 55 nm down to the point of fission [34, 48, 50].

Modified ESCRT-III (C-terminal deletions without MIM) or ESCRT-III fusion proteins as well as catalytic inactive VPS4A or B act in a dominant negative way and block HIV-1 budding upon over expression [17-19, 28, 51].

SiRNA depletion experiments suggested that HIV-1 budding requires in principle only one CHMP4 and one CHMP2 isoform to facilitate egress, whereas CHMP4 filament assembly provides a platform for downstream CHMP2 recruitment [52]. However, CHMP3 cooperates with CHMP2A to increase budding efficiency substantially, while CHMP2B acts independent of CHMP3 [53].

Live cell imaging of budding sites showed that Gag recruitment to the plasma membrane and VLP formation is on average completed within ∼10 min [54]. Alix and Tsg101 (ESCRT-I) progressively accumulate with Gag [55] and once Gag recruitment is terminated ESCRT-III proteins and VPS4 transiently appear at the budding site followed by virus release [55, 56]. Gag co-localizing ESCRT-III clusters showed closed, circular structures with an average size of 45 to 60 nm [57]. Recruitment is sequential with CHMP4B arriving before CHMP2A followed by VPS4. This was suggested to constrict the neck and in case scission does not occur within minutes after ESCRT-III remodeling and disassembly, ESCRT-III and VPS4 are recruited again to the same budding site [58]. The residence time of ESCRTs at the budding site is very short, a few minutes or less [59] and only 1 to 3 % of total budding sites per cell exhibited ESCRT-III and Gag co-localization [57].

In order to facilitate imaging, which is challenged by to the transient nature of ESCRT-III function, particularly in a context where cell physiology is not excessively compromised, we have developed a drug-inducible transient inhibitory ESCRT-III system. We find that inducible CHMP3-NS3-FP had no effect on VLP release in contrast to CHMP3-YFP fusions, but increased the inhibitory effect in combination with CHMP2A-NS3-FP. Furthermore, we demonstrate that inducible CHMP4B-NS3-FP fusions and CHMP2A-NS3-FP fusion proteins can be fine-tuned to reduce Gag-VLP release and temporally increase the number of budding sites revealing Gag-ESCRT co-localization.

## 2 Materials and Methods

### DNA constructs

Homo sapiens CHMP2A, CHMP3, and CHMP4B were fused in-frame with an NS3 cleavage site (wild type or mutated) and the NS3 Hepatitis C protease. They were also tagged with a Flag tag and with either mNeonGreen or mTurquoise (referred to as green and blue respectively or collectively called FP) and subsequently cloned into pCDNA3.1 (for protein sequences, please refer to supplemental Table S1). The mNeonGreen and mTurquoise constructs have been previously described [60, 61].Vps4B R253A, NS3, CHMP2A, CHMP3, and CHMP4B were synthesized (Thermo Scientific). Other plasmids used in this study were previously reported, including pCG-GagRevInd-7ires-puro (referred to as Gag)[62, 63], Gag-mCherry [1], pcDNA-Vphu (ARP-10076, NIH AIDS Reagent Program), GFP-Vps4A E228Q [64] and GFP-p40Phox [65].

### Cells culture, transfection and immunoblotting assay

HeLa cells (ATCC, CCL-2), HEK-293T cells (ATCC, CRL-3216), and Hela Kyoto cells (Bst2+ or Bst2-)[66] were cultured in high glucose DMEM (Gibco) supplemented with 10% FBS (Gibco) and L-glutamine (2mM, Sigma). They were maintained at 37°C in a humidified incubator with 5% CO2. FreeStyle293F cells (Thermo Scientific) grown in FreeStyle293 Expression Medium (Thermo Scientific), were maintained at 37°C with 8% CO2 on an orbital shaker.

For immunoblotting analysis, cells were seeded at 60% confluence into 100 mm dishes and transfected 24 hours later using the Jetprime technique (Polyplus). The cells were co-transfected with 0.5 μg of Gag, the specified amount of CHMP-NS3-FP, and a total transfected DNA amount of 8 μg. Glecaprevir (Cliniscience) was used at a concentration of 25 μM.

Whole cells and VLPs proteins extracts were prepared as previously described [64]. Western blots were conducted using the following antibodies: Anti-HIV-1 p24 (ARP-3537, NIH AIDS Reagent Program), Anti-HIV-1 Vpu (ARP-969, NIH AIDS Reagent Program), Anti-Bst2 (ARP-11721, NIH AIDS Reagent Program), anti-GFP (#A11122, Invitrogen) and Anti-Flag (#F7625, Sigma-Aldrich). Quantification is based on densitometry comparing Gag detection in the VLP fraction to total Gag within cells. To additionally correct for Gag intensities, densitometry values were normalized for Gag to 1 (by dividing the raw mean values for that measured in Gag).

For live cell imaging, cells were seeded in fibronectin-coated glass-bottomed μ-dishes (Ibidi). HeLa CCL2 cells beyond passage p15 and HeLa Kyoto BST2– cells were transfected with an excess of an untagged version of Gag to prevent the morphological defects associated with particles assembled solely from fluorescently tagged Gag [67]. Cells were co-transfected with 0.4 μg of pGag, 0.1 μg of pGag-mCherry, and either 0.5 μg of GFP-VPS4A_E228Q, 0.5 μg of GFP-VPS4B_R253A, or 0.5 μg of pCHMP-NS3-FP. Fluorescent imaging of live cells was conducted 24 hours after transfection, following a 4-hour treatment with Glecaprevir (25µM, Cliniscience) or an equivalent volume of DMSO. An exception was made in the case of the HeLa CCL2 TIRF video-microscopy experiment, where Glecaprevir incubation was limited to 2h.

For electron microscopy imaging, FreeStyle293F cells were seeded in a 6-well plate (Thermo Fisher Scientific) at a density of 2×105 cells per well. After 24 hours, the cultures were co-transfected with 2 µg of pCHMP-NS3-Flag and 2 µg of pGag using 4 µL of 293Fectin Reagent (Thermo Fisher Scientific). Cells were pelleted and high-pressure frozen as described in [68] using an HPM100 system (Leica Microsystems). After freezing, samples were cryo-substituted in an AFS2 machine (Leica Microsystems), dehydrated, and embedded in anhydrous Araldite resin.

### Image acquisition and analysis

Spinning disc microscopy (Cell imaging platform of the IBS) of Gag-mCherry, GFP-p40Phox, CHMP-NS3-blue and CHMP-NS3-green was performed using an Olympus IX81 inverted microscope equipped with a 60X NA1.42 objective (Olympus PlanAPON60X), and CSU-X1 confocal head (Yokogawa). Excitation lasers source (iLaunch, GATACA system) was used for excitation at the suitable wavelengths and power settings. Images were collected employing the Metamorph software (Molecular Devices), via the adapted emission filters set, using a 16b/pixel 512x512 EMCCD (iXon Ultra, Andor).

TIRF video-microscopy was conducted using an inverted microscope (iMIC 2.0, Till-Photonics - FEI) equipped with an alpha-Plan-Apochromat 63x/1.46 objective lens (Zeiss). Image acquisition was performed with an iXon U897 EMCCD camera (Andor). Cells were maintained at 37°C in a 5% CO_2_ environment in a controlled chamber (Ibidi). Time-lapse movies were recorded over a 10-minute duration, with images taken at intervals of 450 ms.

Electron Microscopy: Ultrathin sections were observed using a FEI G2 Tecnai transmission electron microscope (TEM) equipped with an Orius SC1000 CCD camera.

### Single Particle tracking

The open-source Icy software was used to semi-automatically track a large number of individual Gag-mCherry puncta[69]. The procedure included defining the region of interest at the cell’s leading edge, where Gag-mCherry spots were not densely distributed. Gag-mCherry particles were detected with the spot detector plug-in, and their tracks trajectories were subsequently determined using the spot tracking plug-in [69-71]. Manually, tracks at the edge of the region of interest and short tracks (≤ 2) were excluded. Subsequently, the track manager tool was employed to calculate velocities and tracking durations for each particle. Statistical analysis was conducted using GraphPad Prism 9 software.

## 3 Results

To study the ESCRT-III fission machinery *in situ*, within the cellular context at HIV-1 budding site, we developed an assay to transiently enhance ESCRT-III lifetime at budding sites. To this end, CHMP2A, CHMP3 and CHMP4B (referred to collectively as CHMP) proteins were fused to the hepatitis C virus protease NS3 (NS3) via a short linker containing the NS3 cleavage site, which permits auto-cleavage (see Table S1). Treatment of cells with the NS3 inhibitor Glecaprevir induces the temporal accumulation of full-length CHMP-NS3-FP proteins. This accumulation potentially exerts a dominant-negative effect on ESCRT-III function, as previously reported when a heterologous protein is fused to CHMP proteins (Figure 1a) [17, 18].

**Figure 1:**
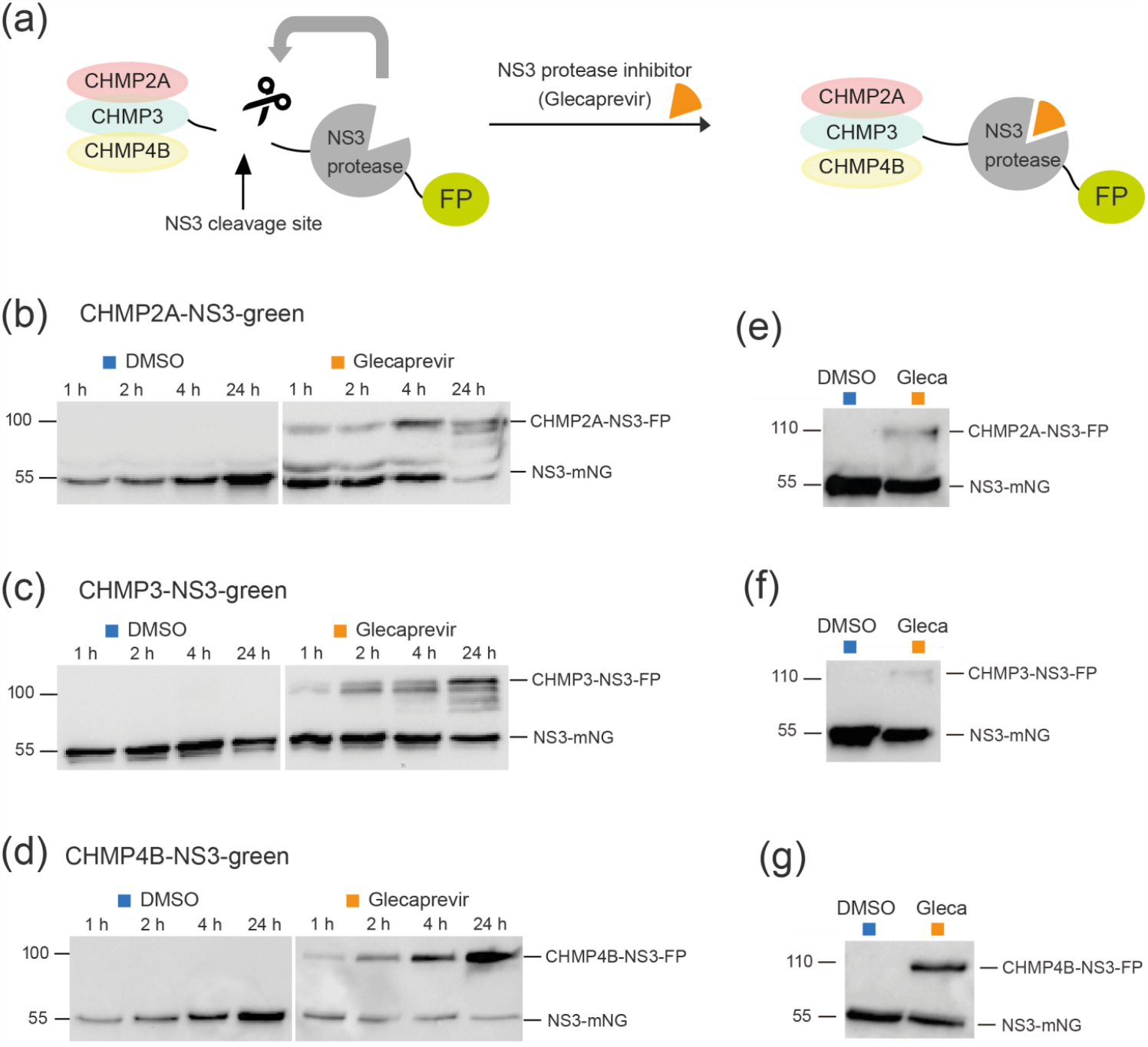
Complete CHMP-NS3-FP proteins are stably expressed over time. **(a)** Schematic illustrating our method for transient expression of ESCRT-III fused to NS3 protease and a fluorescent protein using a drug. ESCRT-III proteins have been fused to the NS3 cleavage site, NS3 protease, and fluorescent protein. Cells transfected with this construct express wild-type ESCRT-III proteins. Upon addition of the drug, a full-length fusion protein accumulates. For imaging and immunoblotting detection, fluorescent proteins (mostly mNeonGreen) and a Flag tag were further added to the NS3 C-terminus, creating CHMP4B/2A/3-NS3-FP-Flag constructs (hereafter collectively referred to as CHMP-NS3-FP) ; **(b – d)** Representative immunoblot experiments were performed using whole cell extracts from HEK293 cells transfected with CHMP2A-NS3-green (2μg) (**b**), CHMP3-NS3-green (2μg) **(c)**, or CHMP4B-NS3-green (1μg) (**d**) and treated with DMSO or Glecaprevir for the indicated duration. **(e – g)** Pulse-chase experiment. HEK293 cells transfected with CHMP2A-NS3-green (**e**), CHMP3-NS3-green (**f)**, or CHMP4B-NS3-green **(g)** were treated with DMSO or Glecaprevir for 4 hours. Subsequently, the medium was washed away and replaced with fresh medium, and whole cell extracts were obtained 24 hours later.

### 3.1 Expression and Auto-cleavage of CHMP-NS3-FP

Immunoblotting experiments were conducted to assess protein expression of CHMP-NS3-FP constructs in HEK-293 cells. Control cells treated with DMSO exhibited a time-dependent accumulation of cleaved NS3-FP proteins after transfection with CHMP2A-NS3-FP (Figure 1b), CHMP3-NS3-FP (Figure 1c, left panels), and CHMP4B-NS3-FP (Figure 1b). Glecaprevir treatment leads to the temporal accumulation of non-cleaved full-length CHMP-NS3-FP proteins as early as one hour (Figures 1b-d, right panels), and longer incubation times led to an increased presence of uncleaved CHMP-NS3-FP proteins. Notably, CHMP4B-NS3-FP showed the highest accumulation of uncleaved fusion proteins upon Glecaprevir treatment, while CHMP2A-NS3-FP reached its maximum after 4 hours and CHMP3-NS3-FP being least efficient in the inhibition of auto-cleavage (Figures 1b-d).

In order to assess the stability of the CHMP-NS3-FP proteins over time, a pulse-chase experiment was conducted. Notably, an increase in the amount of NS3-FP protein was observed, suggesting that newly expressed CHMP2A-NS3-FP, CHMP3-NS3-FP, and CHMP4B-NS3-FP proteins are auto-cleaved (Figures 1e-g). Remarkably, the quantity of full-length CHMP2A-NS3-FP, CHMP3-NS3-FP, and CHMP4B-NS3-FP is not equal and corresponds to the efficiency of fusion protein generation upon Glecaprevir treatment.

We conclude that the addition of Glecaprevir leads to a rapid and stable accumulation of full-length CHMP-NS3-FP proteins over time.

### 3.2 Cellular localization of CHMP-NS3-FP proteins

We next analyzed the localization of transiently expressed CHMP-NS3-FP proteins in HeLa CCL2 cells. In the DMSO group, transfection with CHMP-NS3-FP constructs resulted in a diffuse staining pattern, corresponding to the expression of cleaved NS3-FP (Figure 2a – c). In contrast, after a 4-hour Glecaprevir treatment, while CHMP3-NS3-FP exhibits a diffuse cytosolic staining with few puncta at perinuclear sites and plasma membrane, CHMP4B-NS3-blue and CHMP2A-NS3-blue show some puncta at the plasma membrane and a more pronounced accumulation at perinuclear sites (Figure 2a). To determine the nature of the perinuclear staining, we co-transfected the CHMPs-NS3-blue constructs with the GFP-p40Phox plasmid, which recognizes PtdIns(3)P-enriched early endosomes [65]. We observe a partial co-localization of uncleaved CHMP2A-NS3-blue, CHMP3-NS3-blue or CHMP4B-NS3-blue proteins and GFP-p40Phox following Glecaprevir treatment (Figure 2b, right panels).

**Figure 2:**
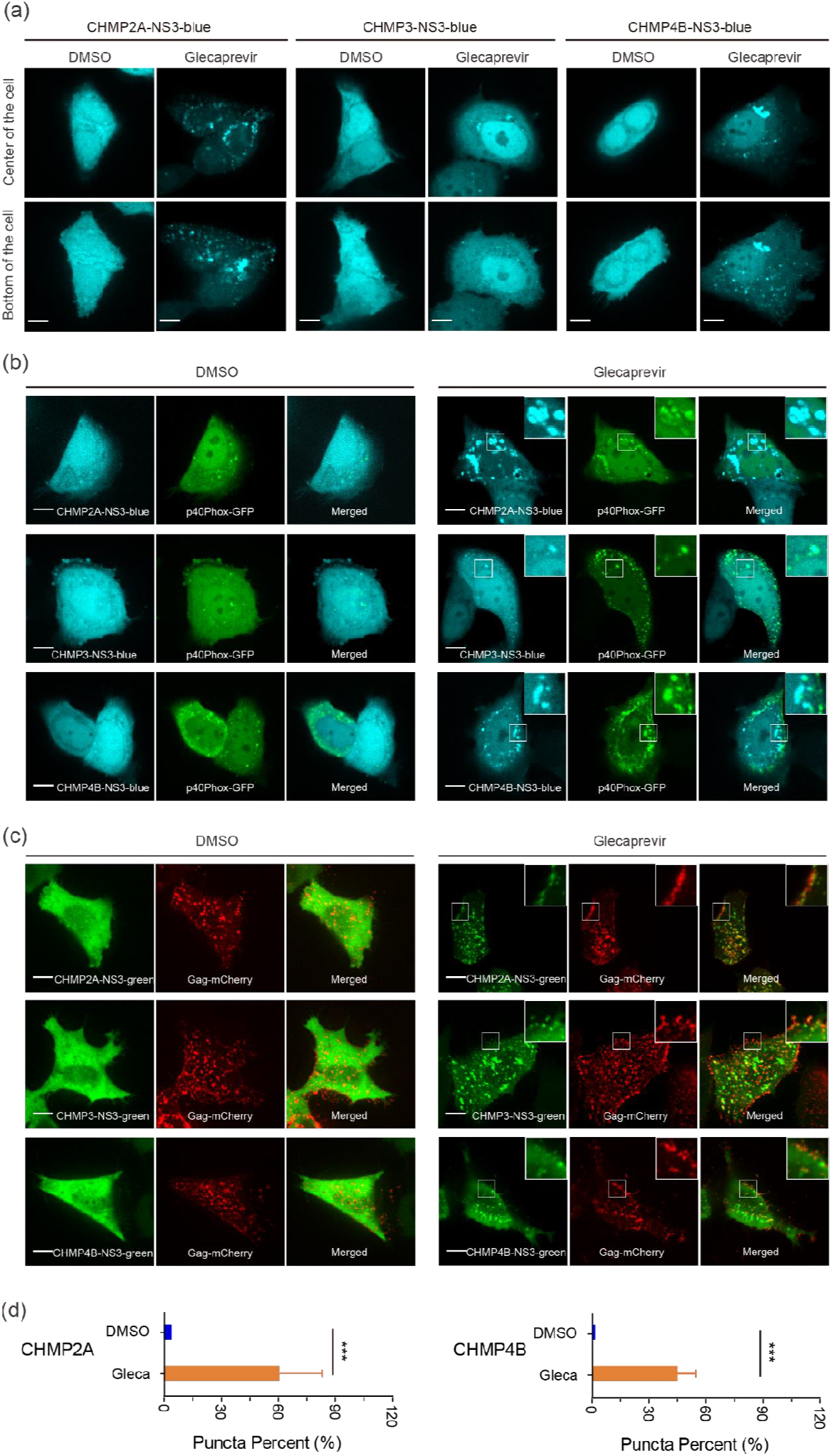
*In vivo* localization of CHMPs-NS3-FP proteins. (**a**) Distribution on CHMP-NS3-FP in HeLA CCL2 cells : Cells transfected with CHMP2A-NS3-blue (left panels), CHMP3-NS3-blue (middle panels) or CHMP4B-NS3-blue (right panels) were treated 4 hours with DMSO or Glecaprevir as indicated. **(b)** CHMPs-NS3-blue accumulate in endosomes : Cells co-transfected with GFP-p40Phox and CHMP2A-NS3-blue (upper panels), CHMP3-NS3-blue (middle panels) or CHMP4B-NS3-blue (lower panels) were treated 4 hours with DMSO or Glecaprevir as indicated. **(c)** Colocalization of CHMP-NS3-green proteins with Gag-mCherry: Cells co-transfected with Gag, Gag-mCherry and CHMP2A-NS3-green (upper panels), CHMP3-NS3-green (middle panels) or CHMP4B-NS3-green (lower panels) were treated 4 hours with DMSO or Glecaprevir as indicated. Scale bars are 10μm. **(d)** Quantification of Gag-mCherry spots proportion co-localizing with CHMP2A-NS3-green or CHMP4B-NS3-green (mean ± SD, n>230 spots for each condition from 3 cells). The P value was calculated using a Mann–Whitney test.

To determine the localization of CHMP-NS3-FP proteins in the context of HIV-1 Gag VLP budding, we co-transfected the CHMP-NS3-FP constructs with Rev-independent HIV-1 Gag (hereafter called Gag) [63]. This showed that DMSO treated cells display cleaved NS3-FP proteins distributed throughout the cytosol. Upon Glecaprevir treatment, CHMP2A-NS3-green, CHMP3-NS3-green and CHMP4B-NS3-green proteins co-localized with Gag-mCherry at the plasma membrane (Figure 2c, right panels). An average of 40% and 60% of Gag-mCherry spots co-localize with CHMP4B-NS3-green and CHMP2A-NS3-green respectively (Figure 2d).

We conclude that Glecaprevir administration permits imaging of Gag co-localization with CHMP2A-NS3-green, CHMP3-NS3-green and CHMP4B-NS3-green at the plasma membrane, indicative of a prolonged half-life of uncleaved CHMP-NS3-FP proteins at HIV-1 Gag budding sites.

### 3.3 Partial Inhibition of Gag VLP Release by CHMP-NS3-FP Proteins

We evaluated Gag–in released VLPs and in Whole Cell Extract (WCE) using immunoblotting in HEK-293 cells. As controls, we included uncleavable CHMP-mutNS3-FP proteins without NS3 protease cleavage site and catalytic inactive dominant negative GFP-VPS4A E228Q [14](Table S1). As expected, Gag co-transfection with GFP-VPS4A E228Q, CHMP2A-mutNS3-FP, or CHMP4B-mutNS3-FP resulted in a significant impairment of HIV-1 Gag VLP discharge, to the extent that emitted VLPs were undetectable compared to wild type Gag VLP release (Figure 3a – c). We also tested the VPS4B R253A mutant that revealed a slower kinetics in disassembling ESCRT-III CHMP2A-CHMP3 helical polymers *in vitro* [64] and demonstrated a reduction of VLP release (Figure 3f). In DMSO-treated cells, VLP emission was minimally affected when CHMP3-NS3-FP or CHMP4B-NS3-FP were expressed while CHMP2A-NS3-FP expression led to a 20% decrease in VLP release (Figures 3a–c).

**Figure 3.**
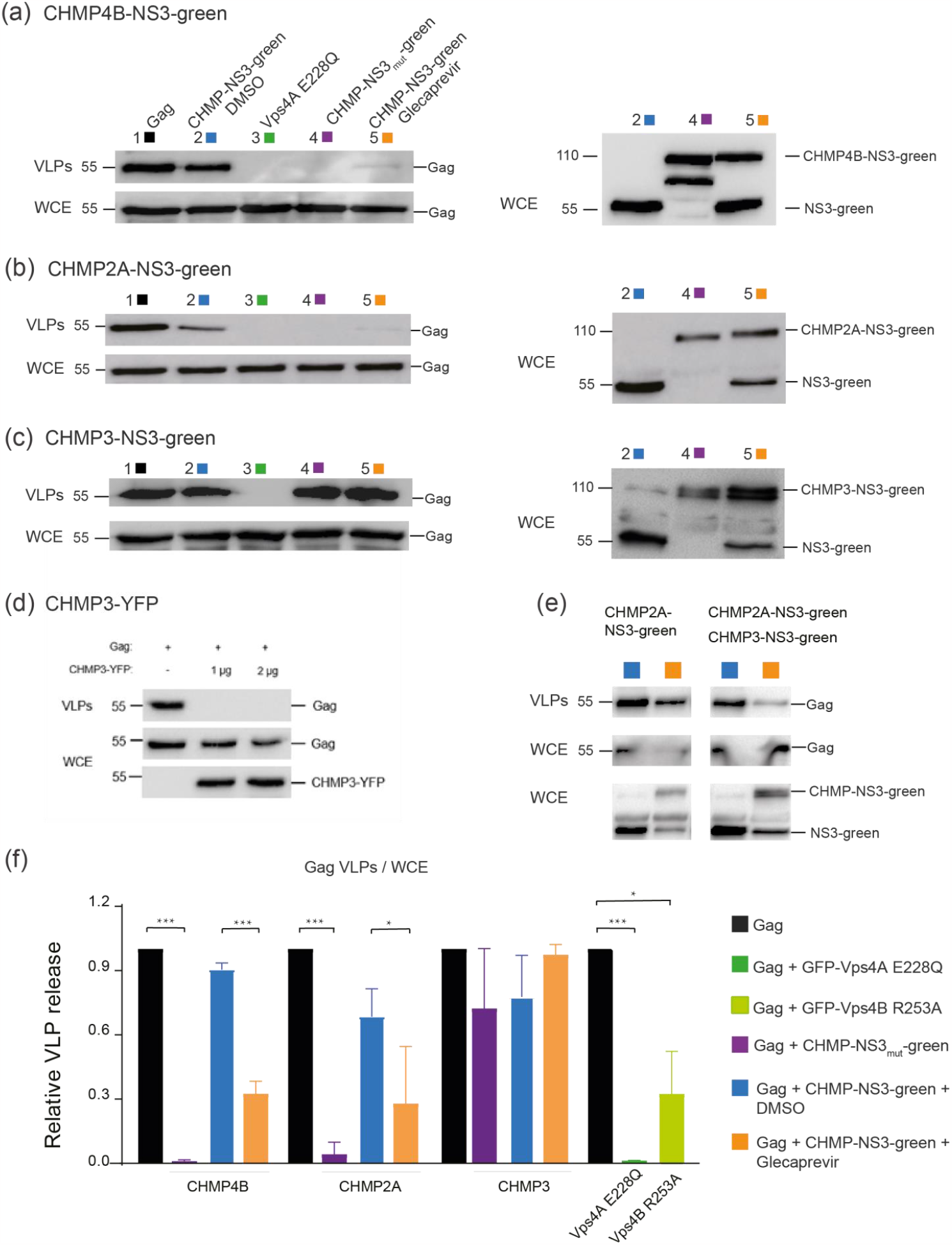
Inhibition of VLP Release by CHMP4B-NS3-green, CHMP2A-NS3-green, and CHMP3-NS3-green: Immunoblot Analysis. **(a-c)** Inhibition of VLP Release in cells co-transfected by Gag (0.5μg), GFP-Vps4A E228Q (1μg), and CHMP4B-NS3-green (1μg) **(a)**, CHMP2A-NS3-green (4μg) **(b)**, and CHMP3-NS3-green (4μg) **(c)**. Representative immunoblot experiment depicting the following: Left panels (i) upper line: HIV-1 VLP pellet, (ii) second line: Gag HIV-1 cellular expression (WCE: whole cell extract), and right panels: CHMPs-NS3-green cellular expression. Hek293 cells were transfected with the following: Column 1: Gag, Column 2: Gag and CHMPs-NS3-green treated with DMSO for 2 hours, Column 3: Gag and GFP-Vps4A E228Q, Column 4: Gag and CHMPs-mut-NS3-green, Column 5: Gag and CHMPs-NS3-green treated with Glecaprevir for 4 hours. **(d)** Representative immunoblot depicting Hek293 cells transfected with Gag and CHMP3-YFP. The upper lines represent HIV-1 VLPs released, the second lines indicate HIV-1 cellular expression (WCE: whole cell extract), and the third lines depict CHMP3-YFP cellular expression. **(e)** Synergy between CHMP3-NS3-green and CHMP2A-NS3-green. To enhance sensitivity the amount of transfected CHMP2A-NS3-green was reduced to 2μg allowing for a slight impairment of VLP release. Co-transfection of Gag (0.5 μg), CHMP2A-NS3-green (2 μg) and CHMP3-NS3-green (2μg) clearly enhance the VLP release inhibition. Representative immunoblot depicting Hek293 cells transfected with left panels: Gag and CHMP2A-NS3-green, right panels: Gag, CHMP2A-NS3-green, and CHMP3-NS3-green, treated with DMSO or Glecaprevir as indicated. The upper lines represent released HIV-1 VLPs, the second lines indicate HIV-1 cellular expression, and the third lines depict CHMPs-NS3-green cellular expression. **(f)** VLP release of the tested constructs analyzed by Western blot. Data is presented for Gag, GFP-Vps4A E228Q, GFP-Vps4B R253A, CHMP4B-NS3-green, CHMP2A-NS3-green, CHMP3-NS3-green, as indicated (mean ± SD, n=3 for each condition). The P value was calculated using a Mann–Whitney test. HIV-1 Gag proteins were detected using anti-p24 antibody, CHMP3-YFP were detected using anti-GFP antibody, while CHMPs-NS3-green were detected using anti-Flag antibody.

Upon Glecaprevir treatment uncleaved CHMP-NS3-FP proteins started to accumulate (Figure 3 a-c right panels) mimicking the effect of the mutation of the NS3 cleavage site in the CHMP-mutNS3-FP proteins (Figure 3 a-c, right panels). Surprisingly, neither CHMP3-mutNS3-FP nor the presence of Glecaprevir in cells transfected with CHMP3-NS3-FP affected HIV-1 Gag VLP release (Figure 3c). However, consistent with previous findings, CHMP3-YFP completely impaired HIV-1 budding [17] (Figure 3d). Although CHMP3 is not strictly required for HIV-1 budding [52], it synergizes with CHMP2A to enhance HIV-1 budding efficiency [53]. Accordingly, we observed a synergistic effect of uncleaved CHMP3-NS3-FP and CHMP2A-NS3-FP on Gag VLP release (Figure 3e). Importantly, expression of CHMP2A-NS3-FP consistently reduced VLP liberation by ∼40%. However, the most pronounced reduction in VLP discharge was observed in cells expressing CHMP4B-NS3-FP, resulting in a decrease of more than 78% without affecting intracellular Gag protein levels (Figures 3a, b, f).

### 3.4 Cell Line-Specific and Dose-Dependent Effects of CHMP-NS3-FP Proteins

Our assay is designed as an imaging tool for studying ESCRT-III proteins prompting us to assess its efficacy in the HeLa CCL2 cell line. Notably, unlike Hek293 cells, HeLa CCL2 cells have been documented to express BST2 tetherin proteins, known for their ability to restrict the release of HIV-1 virions [72, 73]. We thus include a HeLa Kyoto cell line which has been BST2-deleted and stably expresses MKLP1-GFP [66]. As previously reported, HeLa Kyoto BST2+ cells, when transfected with Gag, did not liberate VLPs, while HeLa Kyoto BST2-cells did (Figure S1 a) [73]. Surprisingly, our HeLa CCL2 cells displayed a significant accumulation of free VLPs, indicating a moderate expression of BST2 in this particular HeLa CCL2 cell line. In support of this, immunoblotting revealed reduced BST2 expression in HeLa CCL2 compared to HeLa Kyoto BST2+ cells, with no detectable expression of BST2 in either Hek293 or HeLa Kyoto BST2-cells (Figure S2 b). Furthermore, co-transfection with Vpu, the BST2 inhibiting factor, poorly enhance the accumulation of free VLPs in HeLa CCL2 cells (Figure S1 a). In conclusion, our results suggest that the weak expression of BST2 in the HeLa CCL2 cell line does not efficiently impede the liberation of VLPs.

To assess assay efficiency in HeLa Kyoto BST2-, and Hek293 cell lines, we co-transfected these cells with Gag and varying amounts of CHMP-mutNS3-green or Glecaprevir-treated CHMP-NS3-green constructs, as outlined in Table 1. It is worth noting that in HeLa Kyoto BST2-cells, exposure to Glecaprevir led to a modest reduction in VLP release when using the CHMP4B-NS3-FP and CHMP2A-NS3-FP constructs. Nonetheless, the results unequivocally demonstrate a dose-dependent decrease in HIV-1 VLP release (Table 1). We conclude that budding inhibition efficiency by Glecaprevir-treated CHMP4B-NS3-FP and CHMP2A-NS3-FP proteins is cell line dependent and correlates with expression.

**Table 1.** This table shows the inhibition of HIV-1 VLP budding as determined by immunoblotting assay. Hek293T, HeLa CCL2, or HeLa Kyoto BST2-cells were co-transfected with 0.5 µg of Gag DNA and the indicated amounts of other constructs. n.d. not determined.

| Cell line | HEK293T |  |  |  | HeLa CCL2 |  | HeLa Kyoto BST2- |  |  |
| --- | --- | --- | --- | --- | --- | --- | --- | --- | --- |
| Transfected DNA amount (µg) | 0.5 | 1 | 2 | 4 | 2 | 4 | 1 | 2 | 4 |
| VPS4A_E228Q | n.d. | 100% | 100% | n.d. | n.d. | n.d. | 100% | n.d. | n.d. |
| VPS4B_R253A | 20% | 68% | 79% | n.d. | n.d. | n.d. | 68% | n.d. | n.d. |
| CHMP2A-mutNS3-green | n.d. | 40% | 40% | 100% | n.d. | n.d. | 11% | 18% | 62% |
| CHMP2A-NS3-green<br>Glecaprevir | n.d. | n.d. | n.d. | 41% | n.d. | n.d. | n.d. | n.d. | 6% |
| CHMP3-mutNS3-green | 0% | 0% | 0% | 0% | n.d. | n.d. | n.d. | n.d. | 0% |
| CHMP3-NS3-green<br>Glecaprevir | n.d. | n.d. | n.d. | 0% | n.d. | n.d. | n.d. | n.d. | 0% |
| CHMP4B-mutNS3-green | 64% | 100% | 100% | 100% | 33% | 61% | 20% | 57% | 71% |
| CHMP4B-NS3-green<br>Glecaprevir | n.d. | 78% | n.d. | n.d. | 12% | n.d. | 0% | 13% | 11% |

### 3.5 Quantitative Analysis of HIV-1 Gag VLP Dynamics Reveals Drug-Dependent Prolongation of Gag VLP Retention

We hypothesized that CHMP-NS3-FP’s inhibition of VLP liberation could potentially lead to a delay in VLP release. To precisely measure the timing of VLP release at the cellular level, we conducted thorough TIRF video-microscopy imaging on HeLa CCL2 and HeLa Kyoto BST2-cells. We recorded movies spanning 10 minutes with a temporal resolution of 0.45 seconds per frame. It is important to mention that HeLa Kyoto BST2-cells co-transfected with Gag/Gag-mCherry/CHMP2A-NS3-FP or CHMP4B-NS3-FP and treated with Glecaprevir exhibited a rounded morphology in over 25% of the cells, indicating a potential toxic effect associated with the transfected constructs. Interestingly, this phenomenon was not observed in HeLa CCL2 cells (data not shown). We tracked Gag-mCherry spots and conducted quantitative analyses of their maximum velocity and duration (details in materials and methods). Notably, the distribution of these values appeared consistent across different cells within each experimental condition (Figure S2). Gag-mCherry spot might correspond to (i) VLPs in the process of assembly or assembled but not yet released, (ii) VLPs release from the plasma membrane but tethered by the glycocalix and/or proteins or adhering to the glass surface, and (iii) free VLPs moving in the narrow space between the cell and the coverslip. Free-moving virions have been described to exhibit a high maximum velocity [56, 74]. Consistent with previous findings [56], we observe in both HeLa CCL2 and HeLa Kyoto BST2-cells that the co-expression of VPS4A_E228Q significantly reduced the population of spots with high maximum velocity (> 0.625 μm/s) (Figure S3 a, b),. Furthermore, the expression of both CHMP2A-NS3-FP and CHMP4B-NS3-FP led to a reduction in the number of spots with high maximum velocity (Figure S3 c, d). We conclude that in HeLa CCL2 cells both CHMP2A-NS3-FP and CHMP4B-NS3-FP proteins markedly reduce the number of VLPs released at the individual cell level.

In addition, these findings affirm the accuracy of our tracking analysis.

Next, we analyzed the individual tracking time duration of Gag-mCherry dots. As expected cells transfected with a mixture of Gag-mCherry/Gag and Vps4A_E228Q display a strong increase of spots tracked for the whole movie duration (19.5 + 5.8% compared to 1.5 + 0.9% in control, p<0.0001 unpaired t test Figure 4a). These spots most probably correspond to VLPs assembled but blocked in the process of membrane fission. For CHMP-NS3-FP constructs, Glecaprevir addition enhances the duration time in a specific way. CHMP2A-NS3-FP triggers a 3.4 fold enhancement of Gag-mCherry dots that last for the entire 10 min recording time from 1.4 + 0.7% in DMSO to 4.1 + 1.3% in the Glecaprevir group (p=0.0072 unpaired t test Figure 4c). In contrast, CHMP4B-NS3-FP enhanced the proportion of tracks that have a duration greater than one minute from 7.7 ±0.5% in DMSO to 18.7 ±2.8% in the Glecaprevir group (p<0.0001 unpaired t test Figure 4d). Notably, the observed phenotype of CHMP2A-NS3-FP and CHMP4B-NS3-FP fusion enhancing tracks lasting time is significantly improved when examining co-localized spots (Figure 4c, d).

**Figure 4.**
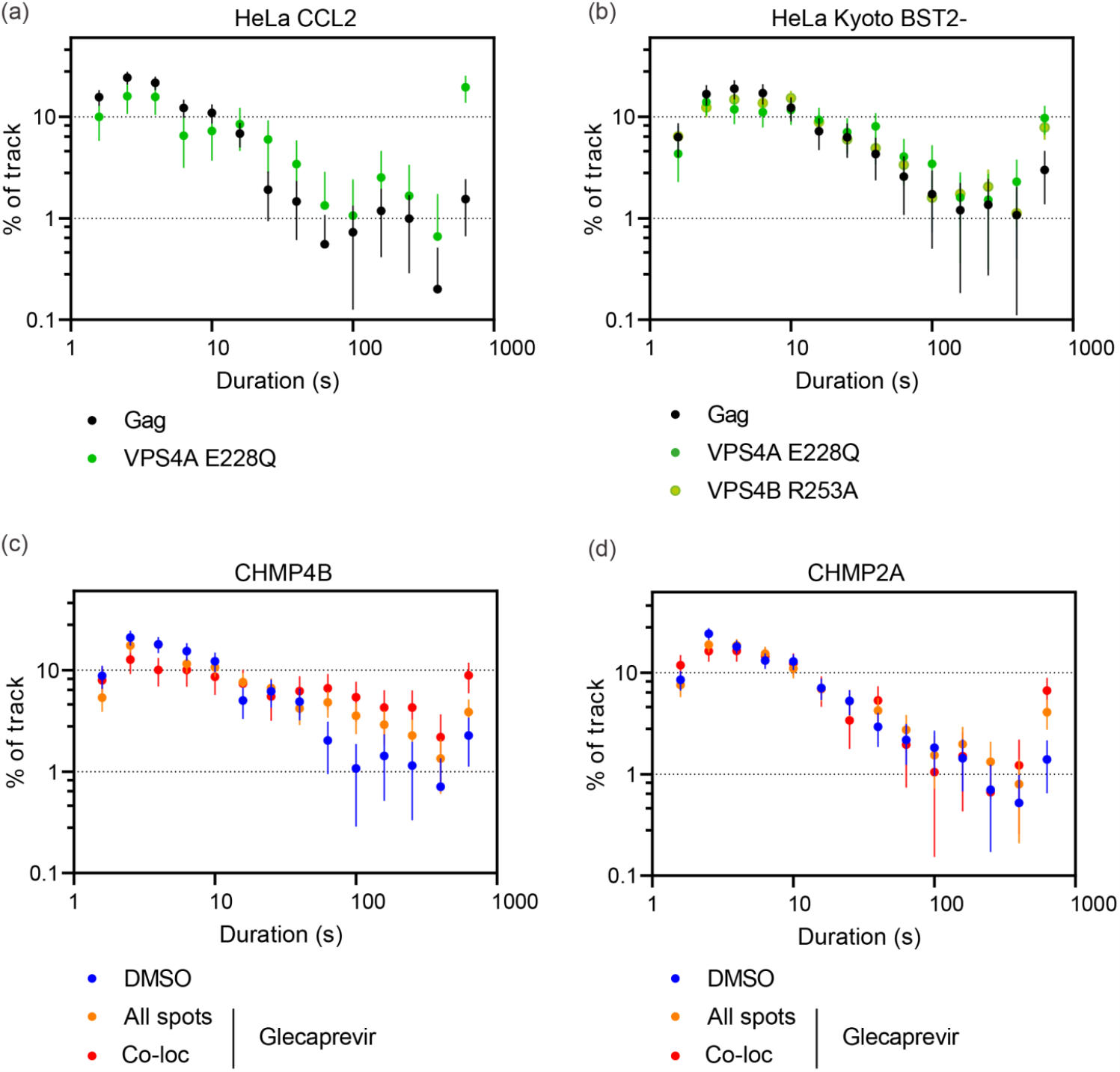
Tracking time duration of Gag-mCherry spots. **(a, c, d)** Frequency distributions of spot tracking time duration in individual HeLa CCL2 cells. The cells were transfected by Gag/Gag-mCherry alone or Gag/Gag-mCherry along with Vps4A E228Q **(a)**, CHMP4B-NS3-green **(c)** and CHMP4B-NS3-green **(d**), and then treated or not by Glecaprevir, as indicated. The error bars represent the standard deviation (n=649; 231; 1874; 1669; 1142; 1527 spots from 4; 5; 8; 8; 8; 8 cells for Gag/Gag-mCherry along with Vps4A E228Q, CHMP2A-NS3-green DMSO, CHMP2A-NS3-green Glecaprevir, CHMP4B-NS3-green DMSO and CHMP4B-NS3-green Glecaprevir respectfully). **(b)** Frequency distribution of spot tracking time in individual HeLa Kyoto Bst2-cells. These cells were transfected with Gag/Gag-mCherry alone, Gag/Gag-mCherry with Vps4A E228Q, or Gag/Gag-mCherry with Vps4B R253A. The error bars represent the standard deviation (n=883; 424; 1744 spots from 9; 6; 9 cells for Gag/Gag-mCherry along with Vps4A E228Q and Vps4B R253A respectfully).

Our findings underscore a drug-dependent prolongation of the life-time of HIV-1 Gag VLPs at cells surface. This effect is particularly pronounced at sites where CHMP4B-NS3-FP and CHMP2A-NS3-FP accumulate.

### 3.6 Electron microscopy reveals Drug induced impairment of Late-stage HIV-1 VLP budding

To gain insight into the structure of VLPs within cells, both in the presence and absence of the drug, we conducted electron microscopy imaging. Specifically, we examined 293FS cells that had been co-transfected with CHMP2A-NS3 or CHMP4B-NS3 along with Gag. These cells were subjected to high-pressure freezing, resin embedding, and subsequent imaging.

In cells treated with DMSO, we observed only a minimal presence of cell-associated VLPs (as depicted in Figure 5a and g). In stark contrast, cells treated with Boceprevir, an NS3 inhibitor, for a duration of 2 hours exhibited numerous particles that remained tethered to the parental cell via membrane stalks. This observation suggests that VLPs had assembled but were unable to undergo release (Figure 5b-e and h-j).

**Figure 5.**
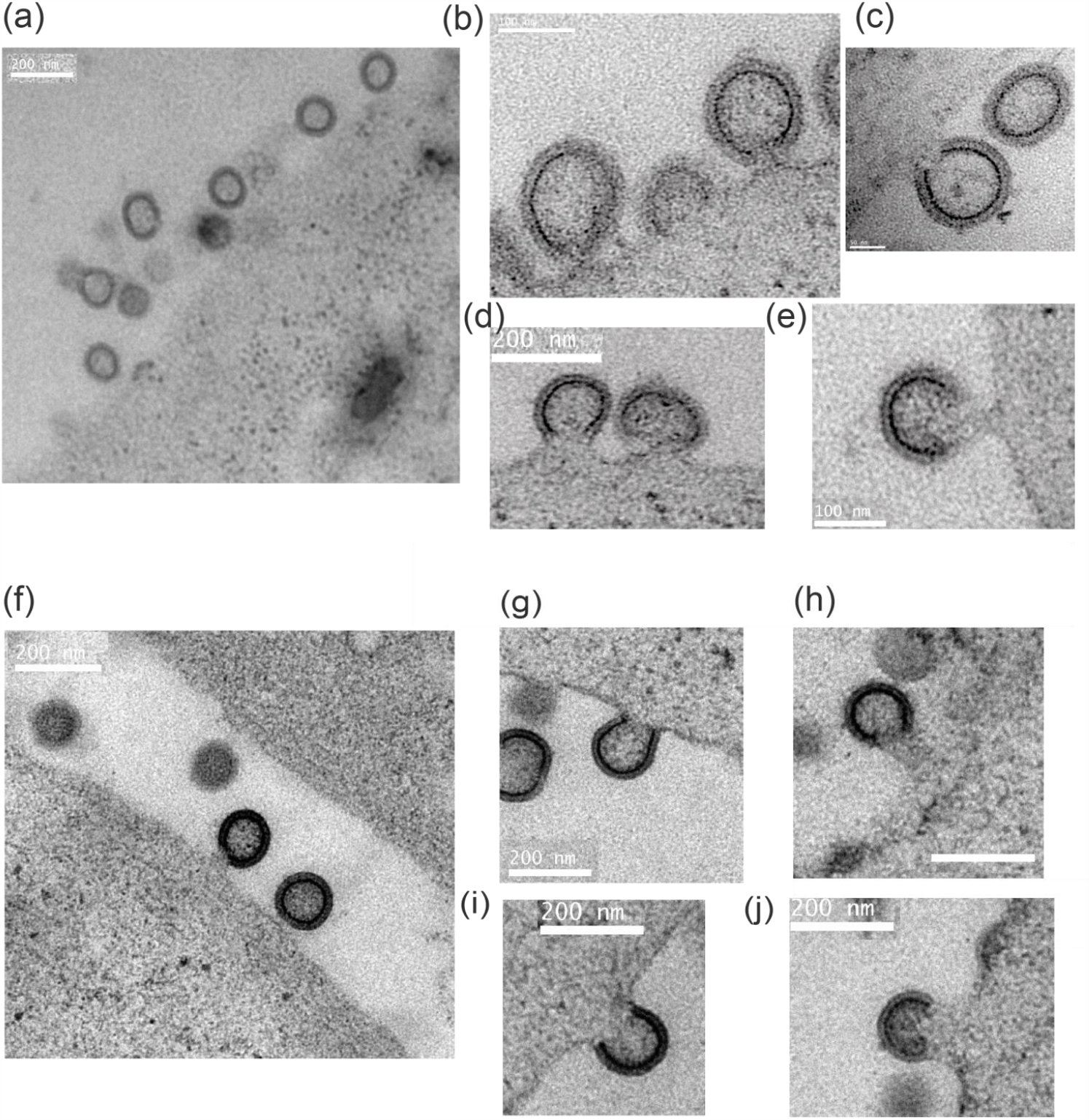
Electron Microscopy Images of 293FS Cells Showing VLP Arrested in Budding at the Plasma Membrane. **(a–e)** 293FS cells transfected with Gag and CHMP2A-NS3 and treated 2 hours with either DMSO **(a)** or Boceprevir **(b– e). (f–j)** 293FS cells transfected with Gag and CHMP4B-NS3 and treated 2 hours with either DMSO **(f)** or Boceprevir **(g– j)**. EM images depict NS3 inhibitor action in arresting VLPs budding at the plasma membrane. Scale bars indicate 50nm **(c)**, 100nm **(b, e)**, and 200 nm **(a, d, f-j)**.

In conclusion, our findings suggest that fusion proteins CHMP2A-NS3 or CHMP4B-NS3 impair the late stage of budding, specifically impeding plasma membrane fission.

## 4 Discussion

A large number of cellular membrane remodeling processes are catalyzed by the ESCRT machinery, which can be efficiently blocked by dominant negative forms of ESCRT-III or VPS4 [22, 75-77]. However, strong dominant negative inhibition affects multiple cellular processes, and the observed effect on a given process may be influenced by blocking many essential cellular functions, not only the targeted pathway. To address this issue, we developed an inducible and transient ESCRT-III inhibition system based on the NS3 hepatitis C protease, described by the Tsien lab [78, 79]. In brief, a cis-acting NS3 protease was fused to the C-terminus of ESCRT-III via a short linker containing the NS3 cleavage site. To track the fusion protein a fluorescent protein (FP) was further genetically fused C-terminally of NS3. The CHMP-NS3-FP fusion proteins undergo auto-cleavage, producing transiently wild-type-like CHMP consistent with previous reports employing cis-acting NS3 [78]. NS3-catalyzed auto-cleavage likely occurs immediately upon expression since no fusion protein can be detected upon one hour of expression. Likewise, inhibition of auto-cleavage with Glecaprevir leads to the transient accumulation of full-length CHMP-NS3-FP fusion proteins detectable within just 30 minutes and maximum levels after 4 hours of treatment for CHMP2A-NS3-green and CHMP3-NS3-green, or 24 hours for CHMP4B-NS3-Green.

C-terminal fusions of CHMP1 have been shown to exert a dominant negative effect on vesicle trafficking [80] and HIV-1 budding can be inhibited with CHMP3-YFP, CHMP3-RFP and CHMP4A/C-RFP fusion proteins [17, 18]. Furthermore, N-terminal ESCRT-III fusion proteins were also reported to be dominant negative [18, 81]. We chose to target the C-terminus because N-terminal fusions of CHMP proteins act differently from C-terminal ESCRT-III fusions. N-terminal fusions likely interfere with membrane interaction due to the N-terminal membrane insertion domains present in most ESCRT-III proteins, which may in turn also affect polymerization on membranes [34, 43, 82]. In contrast, the effect of C-terminal fusions is likely linked to the function of VPS4 which is required for ESCRT-III remodeling and disassembly [25]. Here we show that uncleavable CHMP-NS3-FP fusion proteins containing CHMP2A and CHMP4 effectively impede the release of HIV-1 Gag VLPs. Notably, CHMP3 demonstrates a limited impact on inhibiting VLP discharge, in contrast to the strong inhibitory effects observed with CHMP3-YFP and CHMP3-RFP [17, 18]. We hypothesize that the observed dominant negative effect of C-terminal fusions is due to the overall conformation of the fusion protein, which can sterically interfere with the activity of VPS4. This is consistent with CHMP2B, CHMP3 and CHMP4B tagged with GFP via a long flexible linker (LAP-tag) showing no detectable perturbation of abscission in HeLa cells [37]. We therefore propose that CHMP4B-NS3-FP and CHMP2A-NS3-FP perturb VPS4 activity, which in turn reduces VLP release. In contrast, CHMP3-NS3-FP does not affect VPS4 activity, suggesting that its structure is compatible with VPS4 function. Blocking auto-cleavage of CHMP4B and CHMP2A by NS3 with Glecaprevir does not fully inhibit VLP release and notably blocking auto-cleavage of CHMP3-NS3-FP has no effect on VLP release, but enhances the effect of CHMP2A-NS3-FP, in agreement with the reported synergy of CHMP2A-CHMP3 on HIV-1 budding [53] and CHM2A-CHMP3 heteropolymer formation *in vitro* [34]. Interestingly, a VPS4B mutant that was previously shown to slow down the kinetics of ESCRT-III disassembly *in vitro* [64] shows a similar reduction in VLP release as inhibition of CHMP4B and CHMP2A auto-cleavage. We propose that the inhibited fusion proteins slow down the ESCRT-III/VPS4 machinery leading to a reduced detection of VLPs.

The effect of Glecaprevir, the accumulation of full-length CHMP-NS3-FP proteins, appears to be counterbalanced by protein synthesis, resulting in the continuous presence of cleavage products, NS3-FP and CHMP proteins. The latter is supposed to cooperate with native CHMPs in regular ESCRT-III function. Strikingly, the full-length CHMP-NS3-FP protein accumulation correlates with their dominant negative activity suggesting that the dosage of full-length CHMP-NS3-FP proteins determines the slow down effect of ESCRT-III function.

We demonstrate that the reduction in VLP release is cell line dependent. Hek293 cells exhibit a stronger dominant negative activity of CHMP-NS3-FP, whereas HeLa cells display lower inhibition of VLP release. Notably, HeLa cells have been reported to consistently express BST2, which traps assembled HIV-1 particles at the cell surface and is countered by the HIV-1 accessory protein Vpu [72, 73]. In this study, we observe that BST2 expression is higher in HeLa Kyoto cells compared to HeLa CCL2 cells. Interestingly, in HeLa CCL2 cells, HIV-1 VLP particles are efficiently liberated even in the absence of Vpu, although its presence does enhance the release slightly. These findings are consistent with a certain BST2 threshold concentration being necessary for sequestering HIV-1 particles.

We show further that the NS3-FP fusions affect ESCRT-III localization. Full-length CHMP3-NS3-green is mainly distributed throughout the cytosol, while CHMP2A-NS3-FP and CHMP4B-NS3-FP reveal some punctuate staining at the plasma membrane and at endosomes and/or multi-vesicular endosomes.HIV-1 Gag has been shown to accumulate at the plasma membrane until Gag VLP bud formation that triggers the recruitment of ESCRT-III followed by Vps4A [54, 55, 59]. This recruitment process is of short duration, typically lasting less than two minutes, resulting in a relatively low fraction of HIV-1 budding sites displaying colocalization with ESCRT complexes, estimated to be between 1.5% to 3.4% [57, 58].

In our current study, we provide compelling evidence that non-cleaved CHMP4B-NS3-FP and CHMP2A-NS3-FP proteins exhibit an extended half-life at HIV-1 budding sites, concurrently demonstrating a substantial co-localization of these proteins with Gag-mCherry. In addition, expression of CHMP4B-NS3-FP or CHMP2A-NS3-FP proteins extends the period during which VLP particles could be tracked at the cell surface. We also directly observed an increased number of particles connected to the parental cell by a plasma membrane neck. From these observations, we infer that the accumulation of CHMP4B-NS3-FP and CHMP2A-NS3-FP proteins acts to decelerate the process of plasma membrane fission at HIV-1 budding sites.

In summary, we have set up a novel inducible ESCRT-III system that will be helpful for imaging ESCRT-III at HIV-1 budding sites to obtain more insight into its structure and function. Finally, the system will be useful to study the large plethora of ESCRT-dependent membrane remodeling processes.

## Author Contributions

Conceptualization, W.W.; methodology, W.W. and C.B.; validation, H.W., W.W. and C.B.; Formal Analysis, C.B. and H.W.; investigation, B.G., C.M., C.B., and H.W., resource, J.P.K. and M.P.; data curation, C.B. and H.W.; writing—original draft preparation, W.W. and C.B.; writing—review and editing, C.C., C.B. and W.W..; visualization, C.B. and H.W.; supervision, H.G., C.C., W.W. and C.B.; funding acquisition, W.W. and C.B. All authors have read and agreed to the published version of the manuscript.

## Acknowledgments

We thank Arnaud Echard for the generous gift of cells. HW acknowledges scholarship support from the China Scholarship Council (CSC). WW acknowledges support from the Institut Universitaire de France (IUF). C.B. and W.W. acknowledge access to the platforms of the Grenoble Instruct-ERIC center (IBS and ISBG; UAR 3518 CNRS-CEA-UGA-EMBL) within the Grenoble Partnership for Structural Biology (PSB), with support from FRISBI (ANR-10-INBS-05-02) and GRAL, a project of the University Grenoble Alpes graduate school (Ecoles Universitaires de Recherche) CBH-EUR-GS (ANR-17-EURE-0003). We thank the HIV Reagent Program, Division of AIDS, NIAID, NIH for providing pcDNA-Vphu, Anti-HIV-1 Vpu, Anti-Bst2 and Anti-HIV-1 p24. We thank Guy Schoehn for establishing and managing the IBS electron microscopy platform and for providing training and support. The IBS Electron Microscope facility is supported by the Auvergne Rhône-Alpes Region, the Fonds Feder, the Fondation pour la Recherche Médicale and GIS-IBiSA. IBS acknowledges integration into the Interdisciplinary Research Institute of Grenoble (IRIG, CEA). H.G. was funded by NIH grant R01AI147869.

