## Supplementary material for "An inducible ESCRT-III inhibition tool to control HIV-1 budding": Figure S1

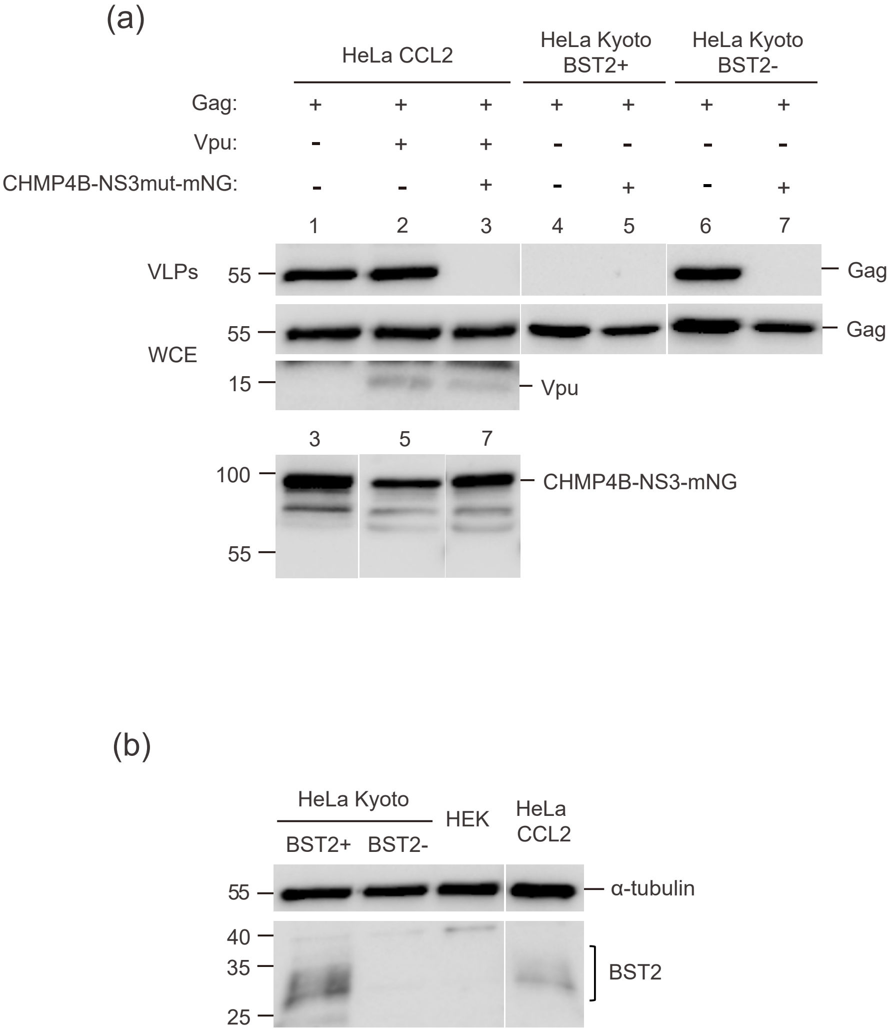


Figure S1: Comparison of Bst2 expression and VLP inhibition in different cell lines.
(a) Representative immunoblot experiment showing HeLa CCL2, HeLa Kyoto Bst2+, or HeLa Kyoto Bst2- cells transfected with Gag, Gag and Vpu, or Gag and CHMP4B-mutNS3-green, as indicated. The upper line represents HIV-1 VLPs released in the medium, the second line represents HIV-1 cellular expression in whole cell extract, the third line represents Vpu expression in whole cell extract, and the fourth panel shows CHMP4B-mutNS3-green cellular expression.
(b) Analysis of Bst2 expression in Hek293, HeLa CCL2, HeLa Kyoto Bst2+, or HeLa Kyoto Bst2- cells. The upper line shows α-tubulin expression in whole cell extract, while the lower panel displays Bst2 expression in whole cell extract.
