## Supplementary material for "An inducible ESCRT-III inhibition tool to control HIV-1 budding": Figure S2

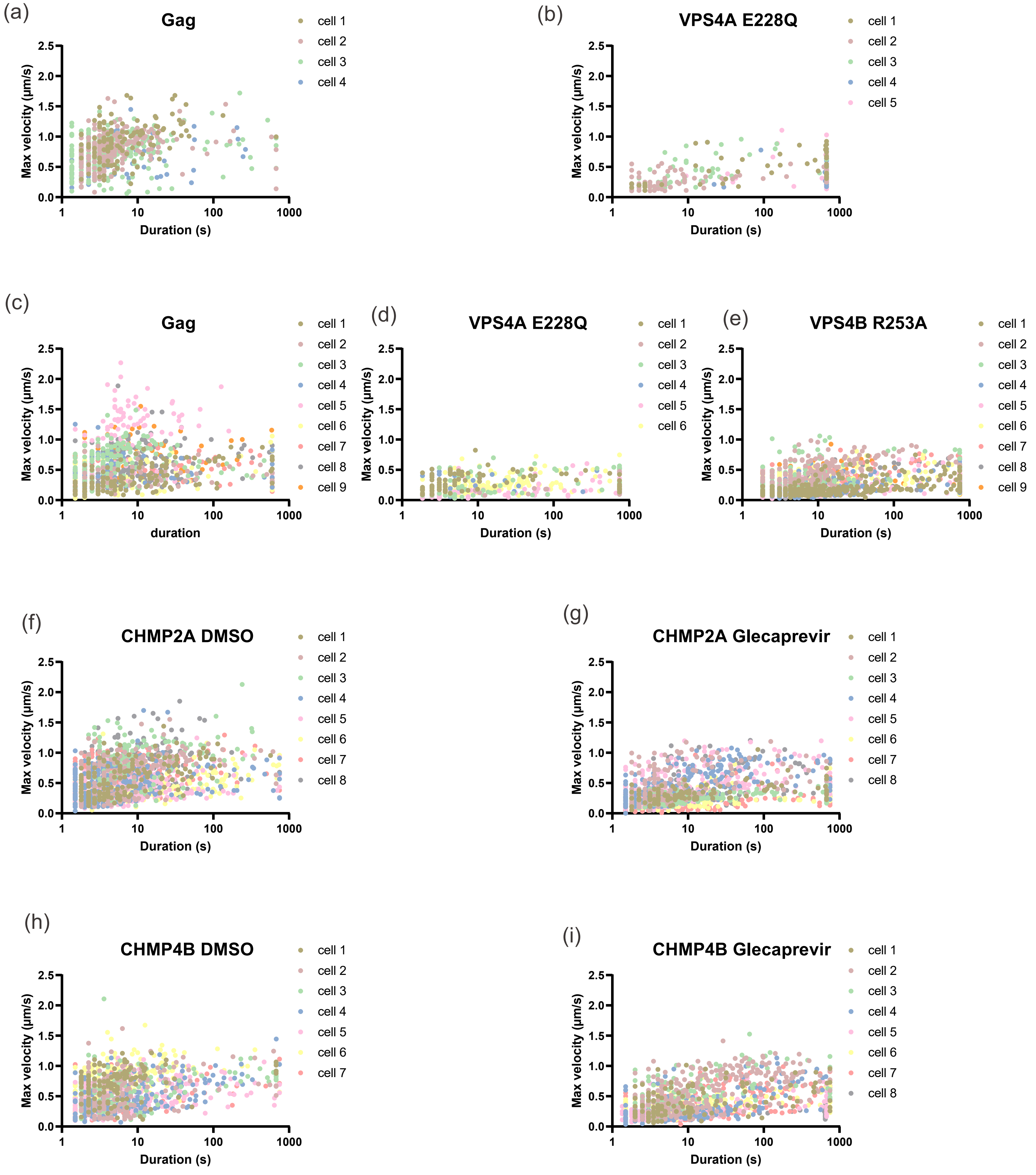


Figure S2: Maximum velocity vs. duration plots of Gag-mCherry spots.
(a, b, f-i) HeLa CCL2 cells were transfected with Gag/Gag-mCherry alone (a) or Gag/Gag-mCherry along with Vps4A E228Q (b), Gag/Gag-mCherry CHMP4B-NS3-green (f, g), or Gag/Gag-mCherry CHMP2A-NS3-green (h, i).
(c-e) HeLa Kyoto BST- cells were transfected with Gag/Gag-mCherry alone (c) Gag/Gag-mCherry along with Vps4A E228Q (d) or Gag/Gag-mCherry along with Vps4B R253A (e). HeLA CCL2 and HeLA Kyoto cells were treated for 2 hours or 4 hours respectivelly with either no treatment, DMSO, or Glecaprevir as indicated.
Each dot represents a Gag-mCherry spot tracked for the indicated duration and maximum velocity.
