## Supplementary material for "An inducible ESCRT-III inhibition tool to control HIV-1 budding": Figure S3

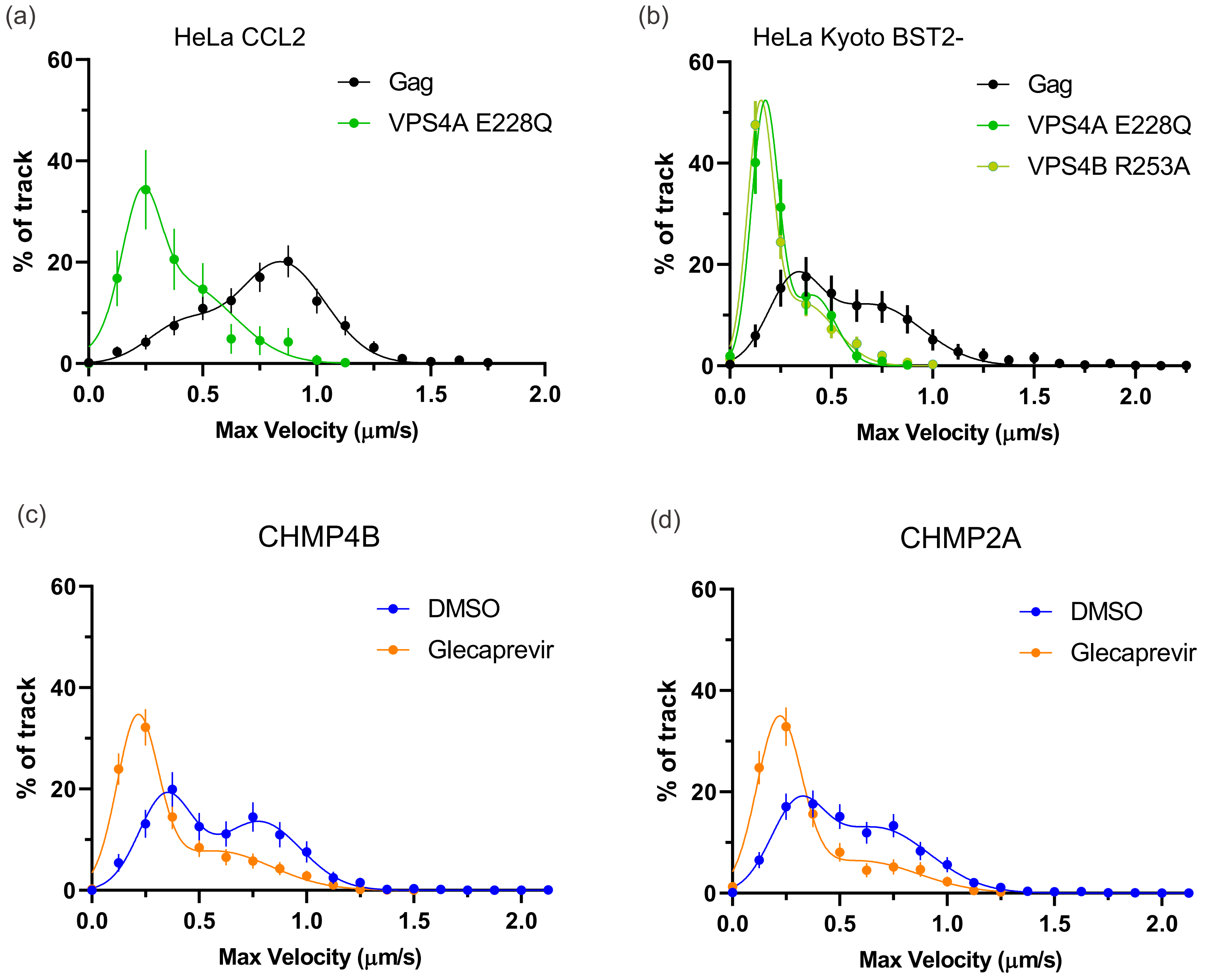


Figure S3: Gag-mCherry Dot maximum velocity Analysis.
(a, c, d) Frequency distributions of spot maximum velocity in individual HeLa CCL2 cells. Cells transfected with Gag/Gag-mCherry alone or Gag/Gag-mCherry along with Vps4A E228Q (a), Gag/Gag-mCherry CHMP4B-NS3-green (c) or Gag/Gag-mCherry CHMP2A-NS3-green (d) were treated 2 hours with either nothing, DMSO or Glecaprevir as indicated. The data was fitted with the best curves, which are a Sum of two Gaussians. The $R^2$ values were 0.98 for Gag, 0.96 for Vps4A E228G, 0.99 for CHMP4B-NS3-green DMSO and CHMP4B-NS3-green Glecaprevir, 0.99 for CHMP2A-NS3-green DMSO, and 0.98 for CHMP2A-NS3-green Glecaprevir.
(b) Frequency distribution of spot maximum velocity in individual HeLa Kyoto Bst2- cells. These cells were transfected with Gag/Gag-mCherry alone, Gag/Gag-mCherry with Vps4A E228Q, or Gag/Gag-mcherry with Vps4B R253A. The best curves were fitted using a Sum of two Gaussians $R^2$ = 0.99.
The error bars represent the standard deviation. spots and cells number are the same as Figure 4.
